# Information Geometry Reconciles Discrete and Continuous Variation in Single-Cell and Spatial Transcriptomic Analysis

**DOI:** 10.64898/2026.02.25.707866

**Authors:** Jinpu Cai, Yuxuan Wang, Yunhao Qiao, Cheng Wang, Ziqi Rong, Luting Zhou, Haoyang Liu, Meng Jiang, Hongbin Shen, Jingyi Jessica Li, Hongyi Xin

**Affiliations:** Global College, Shanghai Jiao Tong University, Shanghai, China; Global Institute of Future Technology, Shanghai Jiao Tong University, Shanghai, China; Bioinformatics Interdepartmental Program, University of California, Los Angeles, CA, USA; Division of Cardiology, Renji Hospital, School of Medicine, Shanghai Jiao Tong University, Shanghai, China; Institute of Image Processing and Pattern Recognition, Shanghai Jiao Tong University, Shanghai, China; Biostatistics Program, Public Health Sciences Division, Fred Hutchinson Cancer Center, Seattle, WA, USA; Department of Biostatistics, University of Washington, Seattle, WA, USA; Department of Statistics and Data Science, University of California, Los Angeles, CA, USA; School of Automation and Intelligent Sensing, Shanghai Jiao Tong University, Shanghai, China

**Keywords:** Single-cell RNA-seq, Information geometry, Cell representation learning, Spatial transcriptomics, Qualitative, Quantitative

## Abstract

Single-cell and spatial transcriptomics provide high-resolution cellular characterization, yet standard analytical approaches remain theoretically misaligned with the probabilistic nature of the data. After UMI normalization, current pipelines rely on Euclidean or log-transformed Euclidean distance for similarity measurement. Both are fundamentally ill-suited to model the multinomial count data. Euclidean distance in normalized space overemphasizes high-variance genes, while log-transformation inverts this bias but at the cost of distorting subtle, continuous expression modulations. Neither approach naturally captures the dual nature of gene expression: both discrete presence/absence transitions and continuous quantitative variation. To overcome these limitations, we introduce GAIA (Geometric Analysis from an Information Aspect), an information-geometric framework for cell representation learning and inter-cell similarity measurement. By anchoring analysis in the true probabilistic model, treating cells as multinomial distributions over genes and projecting cells to a statistical manifold, GAIA organically reconciles both the presence/absence effect and the more continuous expression modulations. Mathematically, GAIA exploits the equivalence between Fisher-Rao distance in multinomial space and geodesic distance on the unit hypersphere, a property that enables both theoretical guarantees and computational efficiency. Experiments in synthetic and real scRNA-seq and spatial transcriptomic datasets demonstrate that GAIA preserves robust and consistent cell-to-cell relationships, delineates biologically nuanced sub-types, mitigates batch effects arising from sequencing depth variation, and eliminates the dependence on knowledge-restricted gene selection for learning meaningful cell representations. Overall, GAIA offers a knowledge-lean, variance-stabilizing framework for analyzing single-cell and spatial transcriptomic data, enhancing discrimination between nuanced cell sub-type and -states.

## 1 Introduction

Recent advances in single-cell and spatial multi-omics technologies have led to increasingly high-resolution characterization of cellular states, enabling the discovery of new cell types, trajectories, and spatial interactions[1]. However, most analytical frameworks still rely on heuristic similarity measures applied directly to normalized gene expression profiles[2, 3]. Typically, cell-to-cell similarities are computed using Euclidean or cosine distances, optionally over log-transformed data, implicitly assuming that the space of gene expression vectors is Euclidean[4]. This assumption is theoretically inconsistent with the underlying probabilistic nature of single-cell sequencing data, which arise from stochastic molecular sampling processes governed by multinomial or related count distributions[5].

Because the Euclidean metric is not derived from the generative model of transcript counts, the relative relationships between cells are highly situational and strongly dependent on the subset of genes used[6]. The geometry implied by these metrics can distort both local and global neighborhood structures in the low-dimensional latent space[7]. As a result, small changes in the selected gene set can lead to substantially different neighborhood structures. Such intricacy requires extensive knowledge and effort in gene selection, in order to produce a meaningful latent landscape. In practice, this often reverts single cell and spatial transcriptomic analysis into a trial-and-error process, increasing the labor effort, reducing the reliability of the analysis, and risking the confinement of the analysis within existing knowledge boundaries[8, 9].

From a probabilistic stand point, Euclidean similarity measurements are dominated by high-expressing genes. Because of the intrinsic stochasticity of the RNA capturing process, high-expressing genes exhibit greater absolute variation and contribute disproportionately to the total distance[10]. A common practical workaround is the log-transformation, which shifts emphasis from absolute expression differences to relative (fold-change) relationships[11]. However, this transformation disproportionately amplifies qualitative differences, such as genes switching from zero to positive expression, introducing strong discontinuities in the geometry of the data space. Consequently, analyses conducted in log-transformed Euclidean space tend to overemphasize presence/absence effects rather than capturing subtler quantitative modulations across genes (Fig 1a).

**Fig 1:**
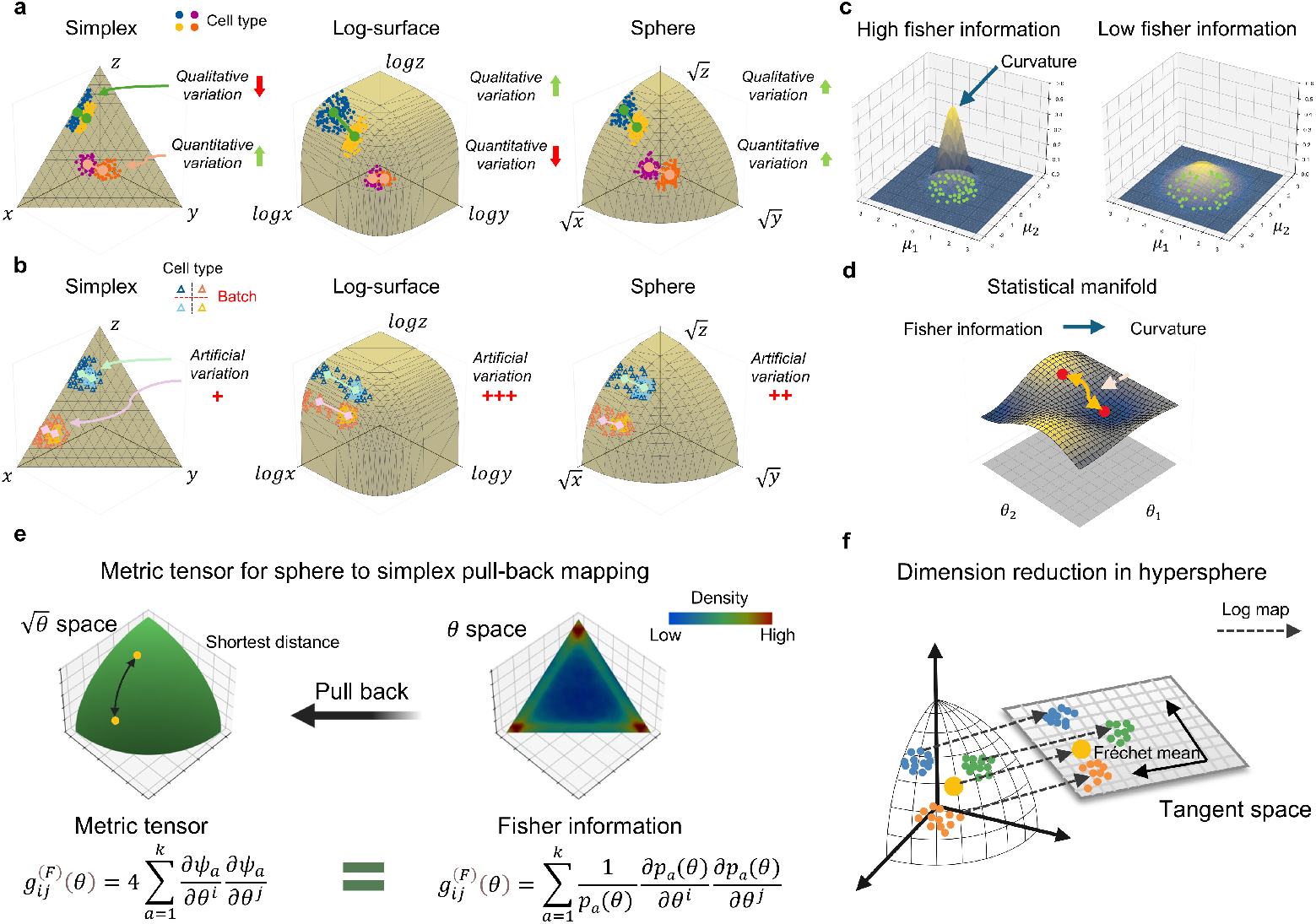
(a) Geometry comparison: Simplex emphasizes quantitative differences, log-surface highlights qualitative variations, and the sphere balances both. (b) Batch effects: Log-normalization introduces artificial variation due to sequencing depth. (c) Fisher information: Measures likelihood curvature, with regions of high and low curvature. (d) Statistical manifold: Parameters form a manifold; distances correspond to geodesics. (e) Pull-back mapping: Under a multinomial model, Fisher information equals geodesic distance on a unit hypersphere after square-root transformation. (f) PCA reduction: Projects the hyper-spherical geometry into lower-dimensional space.

From a geometric perspective, the Euclidean distance between cells on the normalized-and-log-transformed manifold is not geodesic and is highly sensitive to sequencing depth variations (Fig 1b). After log-transformation, the simplex plane stretches into a convex curvature where any Euclidean straight line cuts underneath the surface. Consequently, the Euclidean straight line between two cells is not a geodesic and contains no valid states, as points along it violate the simplex constraint. Thus, the undersurface straight line between two cells on the log-transformed curvature does not represent a viable transition path and its associated Euclidean distance lacks biological meanings. Log transformation also complicates batch integration. When mRNA counts are randomly down-sampled, simulating a shift in sequencing depth, the inter-cell distance experiences an irregular transformation that co-depends on the sequencing depth and gene expression strength: as the relative mean expression a gene approaches zero after down-sampling and gravitating towards becoming a dropout gene, its influence over the inter-cell distance in the log-transformed space grows stronger, disproportionately expanding the dispersion of the cells in its dimension. This irregular distance transformation exacerbates the batch effects by sequencing depth variation and complicates the data integration procedure.

To overcome these limitations, we propose an alternative formulation grounded in *informational geometry* [12]. In our framework GAIA (Geometric Analysis from an Information Aspect), each cell is modeled as a multinomial distribution over genes, representing its normalized expression proportions. The space of such multinomial distributions forms a statistical manifold equipped with a natural Riemannian metric: the Fisher-Rao metric, which measures infinitesimal differences between probability distributions in terms of information distinguishability as Fig 1c,d shows. A key property of this metric is that, for multinomial distributions, under the square-root transformation of the normalized expression proportions, the Fisher-Rao distance between two cells reduces to the angular/arc distance on the positive (1st) quadrant of a unit hypersphere(Fig 1e)[13, 14]. This elegant metric provides both a principled probabilistic foundation and a geometrically interpretable similarity measure which is also computationally efficient.

The GAIA informational geometric representation is knowledge-lean and reconciles the quantitative and qualitative aspects of gene expression variation within a unified metric structure. Quantitative changes correspond to smooth displacements along the manifold, while qualitative transitions correspond to larger geodesic separations reflecting substantial information divergence. Unlike Euclidean distance on a simplex, which steers the measurement towards the quantitative changes of high-expressing genes, nor Euclidean distance on the log-transformed curvature, which is dominated by the qualitative differences of the presence/absence effects, the arc distance on the unit hypersphere after square-root transformation strikes a good balance between preserving quantitative differences while moderately amplifying qualitative shifts (Fig 1a). Most importantly, a geodesic arc on the unit hypersphere define a valid transition path that minimize the integrated information difference between cell states. With the geodesic arc distance, GAIA prevents the underlying statistical manifold to be dominated by either ultra-high-expression housekeeping genes, as in the case of Euclidean distance on a simplex, nor ultra-high-dropout genes, as in the case of Euclidean distance on a log-transformed manifold. Thus, GAIA preserves robust and consistent cell-to-cell distance relations without requiring prudent gene selection, liberating cell representation learning from the tedious, potentially biased and domain-knowledge-intensive gene selection tuning. GAIA also helps mitigating batch effects that stem from sequencing depth variations. Unlike log-transformation, in GAIA, the dispersion growth from the sequencing-depth-induced conversion of a decent-expressing gene to a high-dropout gene is mild (Fig 1b). Thus, sequencing depth variations only introduce mild geometric distortions to the cell distribution in the statistical manifold, naturally improving alignment without sacrificing cell type delineation resolution. In addition to scRNA-seq, GAIA is also instrumental to spatial transcriptomics (ST) analysis. In ST, many genes that are qualitatively distinct at the single-cell level are weakened into merely quantitative or semi-quantitative differences, due to molecule diffusion or oversized spots. By harmonizing qualitative and quantitative differences, GAIA amplifies the nuanced transcriptomic shifts between spots, which is otherwise lost in log-transformation, and improves the spatial domain segmentation accuracy.

## 2 Methods

### 2.1 Probabilistic Framework for ScRNA-seq Data

Single-cell RNA sequencing (scRNA-seq) captures gene expression by sampling a limited number of mRNA molecules from the transcriptome of each cell. Conceptually, each cell can be viewed as a random sample drawn from an underlying gene expression distribution that characterizes its transcriptional state. Thus, the observed count vector for cell *i*, denoted as **x**_*i*_ = (*x*_*i*1_, *x*_*i*2_, …, *x*_*iG*_), represents a finite sampling realization from this distribution. Previous work in [15] has established that single-cell RNA capture can be modeled as a hypergeometric sampling process, in which transcripts are sampled without replacement from the cellular transcriptome. Formally, for a total of *N*_*i*_ transcripts distributed among *G* genes, the probability of observing counts **x**_*i*_ given the true transcript abundances **p**_*i*_ = (*p*_*i*1_, …, *p*_*iG*_) is:

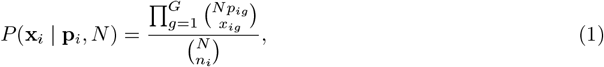

where 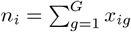 denotes the total number of captured molecules in cell *i*.

When *N* is sufficiently large and the sampling fraction is small (i.e., *n*_*i*_ ≪ *N*), the hypergeometric distribution can be well approximated by a **multinomial distribution x**_*i*_ ∼ Multinomial(*n*_*i*_, **p**_*i*_), with the probability density function as follows:

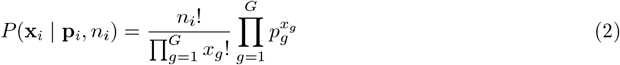

Under this formulation, each cell corresponds to a probability vector **p**_*i*_ lying on the parameter simplex:

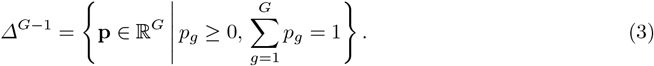

Therefore, the problem of measuring the difference between cells can be naturally reframed as defining a *distance metric between probability distributions* on the simplex, rather than directly comparing raw counts or normalized expression values.

### 2.2 Distance Measurement in the Gene Expression Space

A central task in single-cell analysis is quantifying similarity between cells. Traditionally, distances are computed directly on raw counts or normalized expression vectors. For example, the Euclidean distance between cells *i* and *j* is

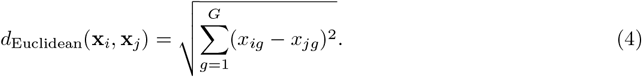

Such approaches, however, are sensitive to sparsity, high dimensionality, and technical noise.

By modeling each cell as a multinomial distribution with probabilities **p**_*i*_, the natural representation becomes

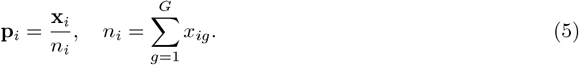

One might consider classical divergence measures between distributions, such as the Kullback-Leibler (KL) divergence

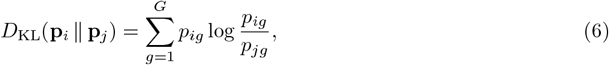

or the Jensen-Shannon (JS) divergence

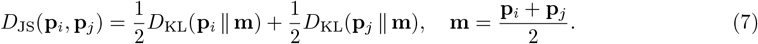

However, KL is asymmetric and unbounded[16], while JS, though symmetric, does not satisfy the triangle inequality, hence not a proper metric distance[17]. From a geometric perspective, neither serves a proper distance metric for cell similarity measurement in the transcriptomic space. Another Wasserstein distance provides an interpretable notion of the “minimal transport cost” between two distributions, it has several limitations in the single-cell context: i) High computational cost, especially for high-dimensional gene spaces; ii) Sensitivity to the choice of cost matrix *c*(*g, h*), which is often arbitrary for genes[18].

While classical distance measures and divergence metrics provide some notion of similarity between cells, they often fail to respect the intrinsic geometry of probability distributions, especially in high-dimensional, sparse single-cell RNA-seq data. This motivates the need for a framework that accounts for the manifold structure of gene expression probabilities, ensuring that distances reflect both local and global variations in the data.

### 2.3 Riemannian Metric Tensor, Fisher-Rao Metric and the Information Geometry

Quantifying distances between cells in single-cell transcriptomics requires a metric that respects the underlying geometry of gene expression distributions. Riemannian geometry, and in particular the Fisher-Rao metric from information geometry, offers a principled approach by defining geodesic-aware distances directly on the statistical manifold of a family of parameterized distributions[19].

A **Riemannian metric** tensor *g*_*p*_ assigns a positive-definite inner product on the tangent space *T*_*p*_ℳ of a smooth manifold ℳ at a coordinate *p* in an Euclidean space:

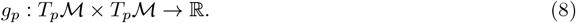

To understand the purpose of the Riemannian metric tensor, one can consider it as a translation between unit vectors from two coordinate systems: the real coordinate in the Euclidean space and a perceived coordinate on the Riemannian manifold. For a coordinate that is defined on the Riemannian manifold, at a given location, the metric tensor provides the means to translate the straight line distance traveled on the manifold, measured in the Riemannian manifold coordinate system, to the aggregated Euclidean distance of the corresponding curve in the real Euclidean space. The real Euclidean length of a tangent vector *v* ∈ *T*_*p*_ ℳ on the Riemannian manifold, defined as the inner product of the vector, can be computed as:

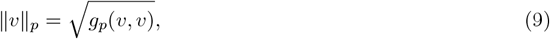

and the geodesic distance between two points *p, q* ∈ ℳ on the non-Euclidean smooth manifold is the length of the shortest geodesic curve connecting them:

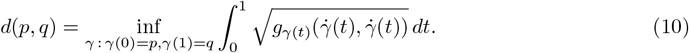

For parametrized distributions, the Fisher information matrix serves as a natural metric tensor that defines a Riemannian geometry in the parameter space. Intuitively, Fisher information matrix, defined as:

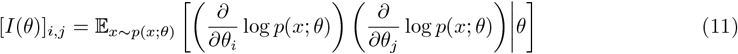

quantifies the variance (if *i* = *j*) or covariance (if *i* ≠ *j*) of the score function (derivative of the log-likelihood function with respect to a parameter *θ*_*i*_ or a parameter pair [*θ*_*i*_, *θ*_*j*_]). Fisher information is always positive-definite. Thus, it can be perceived as defining a curvature of the support curve representing the local geometry of a parametrized distribution in its parameter space. High Fisher information for a parameter(s) indicates that the curvature along the parameter(s) direction is sharp, therefore minute perturbation of the parameter(s) would significantly alter the distribution. Low Fisher information indicates the curvature is blunt and thus the nature of the distribution is less affected by changes in the respected parameter(s).

Given the natural equivalence of the Fisher information matrix and the Riemannian metric tensor, we can harness the Fisher information matrix as defining a mapping between a Riemannian geometry, in which the distributions locates on a Riemannian surface, and the Euclidean parameter space. The distance, known as the **Fisher-Rao distance**, between two distributions *p*(*x*; *θ*_1_) and *p*(*x*; *θ*_2_) with respective parameters can thus be computed as the geodesic distance of the shortest path on the Rie-mannian statistical manifold:

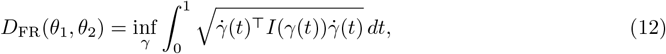

where *γ*(*t*) is a smooth path connecting *θ*_1_ and *θ*_2_. The Fisher-Rao distance is symmetric, invariant under reparametrization, and respects the underlying probabilistic structure, making it well-suited to quantify cell–cell similarity in single-cell RNA-seq data.

The Fisher-Rao distance quantifies how distinguishable two probability distributions are given their coordinates in the parameter space. Intuitively, with a high Fisher Information value defining a sharp curvature, distributions in a local pocket are more different in nature as they are less capable of explaining samples drawn from each other. These distributions are more distant on the Riemannian manifold as they have to overcome a large vertical distance on the statistical manifold despite their proximity in the parameter space. The Fisher-Rao distance, an integral of the shortest curve on the Riemannian manifold, therefore measures the aggregated difficulty of reshaping one distribution into another through gradually updating the parameters.

### 2.4 Square-root Transformation and the Simplex-to-Sphere Pull-back Mapping

The Fisher-Rao distance does not always produce a well-known, regular geometry (e.g., spherical, hyperbolic). Multinomial distribution, among a few others, is an exception where the information geometry is the first quadrant of a hypersphere. To show this, we need to demonstrate that the Fisher information matrix for multinomial distributions is equivalent (amplified by a constant factor) to the metric tensor for a pullback mapping from a simplex to a unit hypersphere, with the coordinates on the sphere being substituted by its squared variant, which is the coordinates on a hyper-simplex. The coordinate substitution could be comprehended as a pullback mapping of the metric tensor on a hyper-sphere space to the Euclidean hyper-simplex system, with square-root transformation.

Let ***θ*** denote the *n*-dimensional Cartesian coordinates of points on a simplex, where |***θ***| _1_ = 1. Let ***ω*** denote the square-root transformed Cartesian coordinates, in which the simplex is transformed into a sphere with |***ω***|_2_ = 1. For a point on the sphere, let the bases ***e***_**1**_, ***e***_**2**_, …, ***e***_***n***_ denote the orthogonal bases of the local tangent space. A vector ***z*** in the local tangent space, represented as 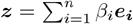 can be converted to a representation in the ***ω*** space as 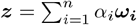 with chain rule, where 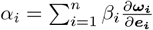. The inner product of a vector ***z*** in the ***ω*** coordinate system can thus be translated to the local tangent space as following:

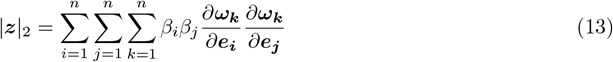

The above equation can be perceived as a translation of L-2 norm between the local tangent space and the ***ω*** coordinate system, as |***z***|_2_ = ***z***⊤***Gz***, where ***G*** is the metric tensor defined as 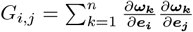. By substituting in 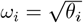 and applying chain rule in reverse 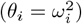, we have:

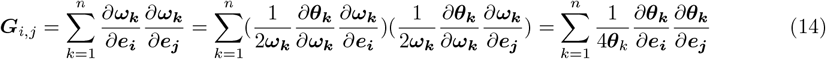

given that 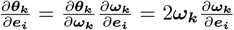. Equation 14 establishes the metric tensor between a simplex and its square-root-transformed unit hypersphere, where the geodesic distance on the equivalent hypersphere manifold between two points ***θ***_1_ and ***θ***_2_ on the hyper-simplex can be computed as:

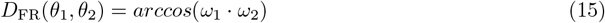

with *θ*_1_ = (*ω*_1_)^2^ and *θ*_2_ = (*ω*_2_)^2^.

For multinomial distributions, the Fisher information matrix takes the same form as the metric tensor for the simplex-to-sphere pullback mapping. To see this, we use an alternative form of the Fisher information matrix

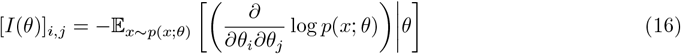

By plugging Equation 2 into Equation 16 and apply chain rule, we have:

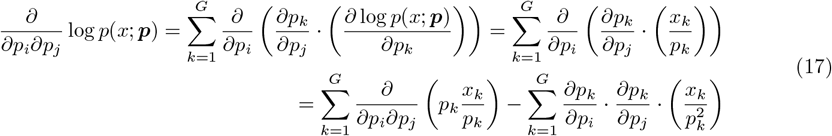

Take 17 into 16, we have:

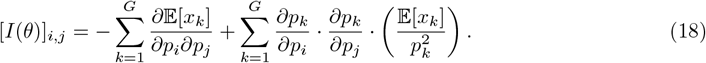

Notice that 𝔼 [*x*_*k*_] = *N* · *p*_*k*_ with *N* being the total RNA molecule count of a cell. Thus Equation 18 becomes

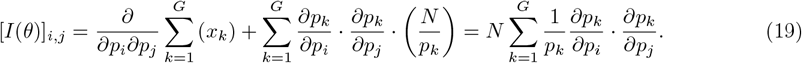

Notice that Equation 19 has the same form as Equation 14 but amplified by a factor of 4*N* . This establishes the geometric equivalence between information distance and arc distance: The Fisher-Rao distance between two multinomial distributions is a constant multiple of the arc distance between ***ω***_*i*_ and ***ω***_*j*_ on the hypersphere, after taking square-root transformation: 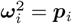 and 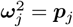.

### 2.5 Spherical Embedding Preserves Transcriptomic Heterogeneity in High Dimensions

An important property of the spherical embedding for multinomial distributions is that it makes feature engineering such as PCA less sensitive to gene selection. Specifically, the spherical embedding preserves the geometry that appears in a low-dimensional informative gene space, after expanding the gene space into higher dimensions by adding uninformative noisy genes. This property of being robust to noisy housekeeping genes is an direct corollary of an even stronger statement, which states that Fisher information, and the spherical embedding for the multinomial distributions, is isometry for every Markov embedding. This is known as the Chentsov theorem[20].

The significance of Chentsov theorem is that it establishes the theoretical basis for Fisher information metric being the most natural geometry to quantify the distance between distributions. It shows that Fisher information is the only isometric Riemannian geometry for all Markov embeddings. Retaining isometry is a crucial property for an effective dimension reduction regime. Assume mRNA molecules from *G* genes are partitioned into *M* biological functions (mRNA but not genes are partitioned). Let 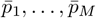 denote the probability of drawing mRNA molecules from the respective biological functions. Let *q*_*m*,1_, …, *q*_*m,G*_ with ∑_*i*_ *q*_*m,i*_ = 1 denote the probability of a mRNA molecule of a biological function *m* belonging to specific genes. Then the probability *p*_1_, … *p*_*G*_ of drawing mRNA from specific genes, could be seen as a compound two-step multinomial sampling: first from 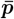 and then from ***q***. We call ***q*** being the Markov embedding which maps one multinomial distribution function space of *M* dimensions 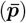 to another multinomial distribution function space of *G* dimensions (***p***). Chentsov shows that Fisher-Rao distance is the only metric where the distance relations between distributions remain the same after any Markov embedding: for two pairs of distributions, *<* ***p***_1_, ***p***_2_ *>* and *<* ***p***_3_, ***p***_4_ *>* where *D*_FR_(***p***_1_, ***p***_2_) = *D*_FR_(***p***_3_, ***p***_4_), then 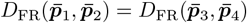 after transformation by any Markov embedding regime ***q***.

The Euclidean distance between the proportions of genes ***p***, for instance, is not isometry. Consider two pair of cells *< c*1, *c*2 *>* and *< c*3, *c*4 *>* in a 3-gene space. Let the ratio between *g*1 : *g*2 = *r*1 in both *c*1 and *c*3, and *g*1 : *g*2 = *r*2 in both *c*2 and *c*4. From the perspective of the dual-gene space *g*1, *g*2, *< c*1, *c*2 *>* and *< c*3, *c*4 *>* has the same Euclidean distance: *d*_[*g*1,*g*2]_(*c*1, *c*2) = *d*_[*g*1,*g*2]_(*c*3, *c*4). However, once we consider the third gene *g*3, the distance equivalence is no longer maintained. Let *c*1, *c*2 has the same, high (but less than 50%) *g*3 proportion; while *c*3, *c*4 has equal, low *g*3 proportion. It is easy to see that *d*_[*g*1,*g*2,*g*3]_(*c*1, *c*2) *< d*_[*g*1,*g*2,*g*3]_(*c*3, *c*4). This means that a constant ratio between *g*1, *g*2 is interpreted differently with respect to different expression strengths of an irrelative gene *g*3, creating an unnecessary attachment between genes. The spherical embedding on the other hand, always maintain the distance equivalence between *< c*1, *c*2 *>* and *< c*3, *c*4 *>* in both 2-gene space and 3-gene space. Therefore the difference between *g*1 : *g*2 = *r*1 and *g*1 : *g*2 = *r*2 remains a constant irrespective of including other irrelevant genes (Fig 2).

**Fig 2:**
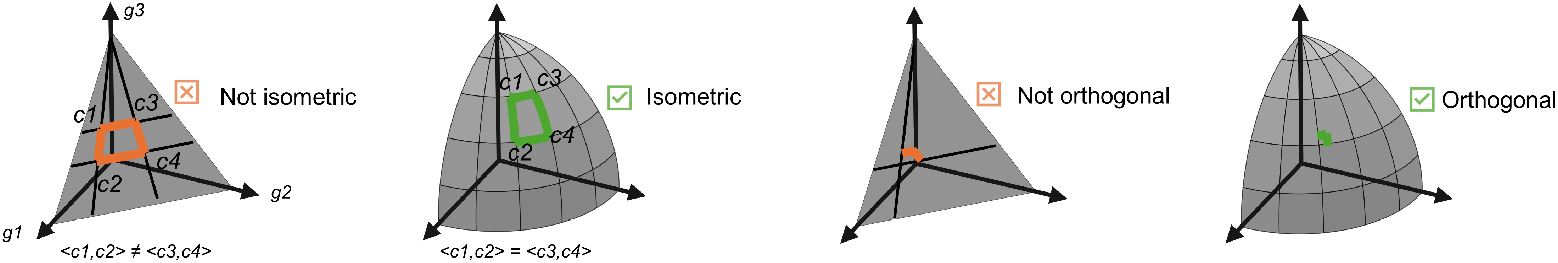
Spherical embedding ensures isometry and orthogonalization of distances.

The disentanglement between genes could also be explained from a geometric perspective. Assume there are two groups of cells in a 3-gene space. The first group *S*1 has the same *g*3 proportion but varies in *g*1 : *g*2 ratios. The second group *S*2 has a constant *g*1 : *g*2 ratio but varies in *g*3 proportions. In the Euclidean simplex, *S*1 and *S*2 are distributed along two straight lines but these two lines are only perpendicular if either cells in *S*2 have *g*3 = 50% or cells in *S*1 have *g*1 : *g*2 = 1. Otherwise, *S*1 and *S*2 are not truly orthogonal and *g*3 becomes entangled with *g*1 : *g*2 ratio. On the Riemannian sphere, in contrast, *S*1 and *S*2 are always on a pair of strictly perpendicular longitude and latitude. Thus, *g*3 is always locally orthogonal to *g*1 : *g*2 ratio shifts at their intersection (a latitude is not a straight line on the spherical surface but is locally orthogonal to any longitude). This geometric orthogonality is key to the establishment of a stable, gene-selection-invariant dimension reduction regime in the spherical manifold. Variance in irrelative uninformative genes will always be orthogonal to meaningful ratio shifts between biological functions and are expelled from the top low-dimensional components.

### 2.6 Harmonization of Qualitative and Quantitative Differences with Square-Root Transformation

Both the log transformation and the square-root transformation are special cases of the Box-Cox transformation family[21]. Formally, the Box-Cox transformation is a parametric family of power transformations:

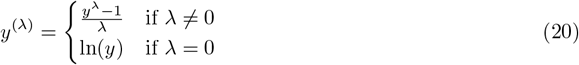

The log and the square-root transformations are special cases where λ = 0 and λ = 0.5 respectively. The Box-Cox transformation is designed to stabilize variance and induce approximate normality in non-normal data. After Box-Cox transformation, the variance is expected to be detached from the mean of the random variable, such that distance measurements would not be biased towards the highest-expressing genes. In Box-Cox transformation, λ characterizes the anticipated relationship between the mean and the variance of a random variable. For distributions where standard deviation scales proportionally to mean, such as members in the Gamma distributions family, log transformation is particularly effective where it converts multiplicative relationships into additive ones. For distributions where variance scales proportionally to mean, such as Poisson, square-root transformation is the most effective. Multinomial distributions, with its mean (𝔼 [*X*_*i*_] = *np*_*i*_) and variance (*V ar*(*X*_*i*_) = *np*_*i*_(1 − *p*_*i*_) = (1 − *p*_*i*_)𝔼 [*X*_*i*_]) being approximately linearly dependent, square-root transformation is therefore the more appropriate transformation.

Imprudent application of log transformation to multinomial observations could lead to a number of adverse effects: 1) amplification of low-count/boundary effects: When *p*_*i*_ is small, log transformation pulls values near zero close to −∞, amplifying the variance of low-expressing genes and introduces distortions to distance measurements. 2) Loss of quantitative information: With an over-emphasis on the multiplicative structure, log transformation assigns equal distances between 1 → 5 and 100 → 500. A difference between 100 → 200, generally considered a significant shift in expression strength, pales in comparison to the minute perturbation of a gene growing from 1 → 5 due to sequencing randomness. 3) Asymmetric variance stabilization across the data range: For a large *p*_*i*_, log transformation significantly compresses the scale, leading to a collapse of the variance. But for a small *p*_*i*_, log transformation could drastically expand the scale, leading to an inverse inflation of the variances for small means. The co-dependence between mean and variance in log transformation is best illustrated by computing the asymptotic variance. Using the delta method by leveraging the first-order Taylor expansion, we have 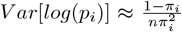, where *π*_*i*_ = 𝔼 [*X*_*i*_] = *np*_*i*_. As *π*_*i*_ → 0, the variance explodes to ∞. In contrast, for square-root transformation, we have 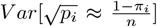, where the variance is significantly more stable across the gene expression strength spectrum.

To systematically evaluate how data transformation affects the balance between qualitative and quantitative variation in single-cell data, we adopted the Box–Cox transformation as a unifying analytical framework. We simulated datasets containing both discrete and continuous expression differences among cell types and quantified their separation using class-center distances. To measure the overall preservation of biologically meaningful variation, we defined a harmonization score: the harmonic mean of qualitative and quantitative distances, which penalizes transformations that overemphasize one type of variation at the expense of the other. This framework allowed us to identify the transformation that most effectively maintains a balanced representation of discrete and continuous transcriptional differences across the expression range. We found that as λ increases, the captured variation gradually shifts from qualitative to quantitative, and the harmonization score reaches its maximum at λ = 0.5, corresponding to the square-root transformation.

While square-root transformation is effective at stabilizing variance for multinomial distributions and does not completely diminish the intrinsic variances for high-expressing genes, it is worth noting that simply regarding the square-root transformed gene proportion space as a Euclidean space is still inappropriate. Euclidean distance, a natural metric for Normal distributions, is fundamentally incompatible with multinomial distributions, despite that square-root transformation reshapes multinomial distributions to be more Normal-like. Fundamentally, the Riemannian arc distance on the square-root-transformed hyper-sphere is the most natural metric for multinomial distributions, which doubles as the Fisher-Rao metric on the probability simplex.

### 2.7 Dimensionality Reduction on the Square-Root Transformed Sphere

After applying the square-root transformation to map multinomially-distributed single-cell data onto the unit sphere, standard linear dimensionality reduction methods such as PCA are no longer directly appropriate, since the data now reside on a curved manifold rather than in Euclidean space. To address this, we employ *Tangent PCA*[22], which respects the spherical geometry while providing an approximate low-dimensional embedding.

Let **x**_1_, …, **x**_*N*_ ∈ 𝕊^*G*−1^ denote the square-root transformed expression profiles of *N* cells on the unit sphere in *G* dimensions. The first step is to estimate the *Fréchet mean* 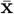 of the points on the sphere:

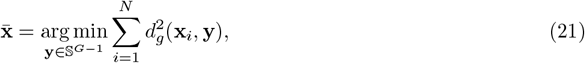

where 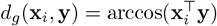 is the geodesic distance on the sphere. Next, each point is mapped to the tangent space at 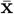 using the logarithm map:

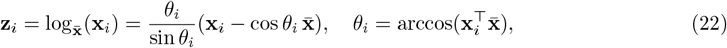

yielding vectors **z**_*i*_ in the (*G* − 1)-dimensional tangent space. Standard PCA can now be applied in this tangent space:

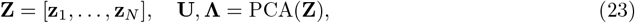

where **U** contains the principal directions and **Λ** the corresponding variances.

The tangent plane 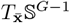 provides a local linearization of the sphere around the Fréchet mean 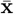. For points **x**_*i*_ close to 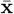, the Euclidean distance in the tangent plane approximates the geodesic distance on the sphere:

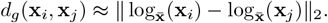

Consequently, performing PCA in the tangent plane yields a local linear approximation to spherical PCA, preserving the principal directions of variation while respecting the sphere’s geometry.

## 3 Results

### 3.1 GAIA Robustly Identifies Stable B Cell Subtypes Across Feature Selections

We applied GAIA to the bone marrow mononuclear cell (BMMC) dataset from [23], and performed clustering using the Leiden algorithm on the GAIA embedding. Using the GAIA framework, we delineated four transcriptionally distinct B cell subtypes (Fig 3a), each characterized by coherent gene expression programs. In contrast, the four B cell sub-types become considerably inter-mixed in the normalization-only and the normalization-and-log-transformation frameworks. Marker genes for each cluster were identified via Wilcoxon rank-sum test performed on normalized count matrices. In GAIA-derived clusters, we observed distinct immunoglobulin expression programs. Specifically, two clusters were enriched for CD27, consistent with a memory B cell phenotype, while others showed differential enrichment of IGKC (*κ* light chain) and IGLC2 (λ light chain), reflecting mutually exclusive light chain usage [24]. These marker genes, together with IGHM and IGHD, are well-documented canonical markers distinguishing B cell maturation stages and functional states, as curated in the CellMarker database [25]. Notably, we observed that these marker genes exhibited both quantitative differences in expression levels and qualitative differences in presence/absence patterns across the four clusters (Appendix A).

**Fig 3:**
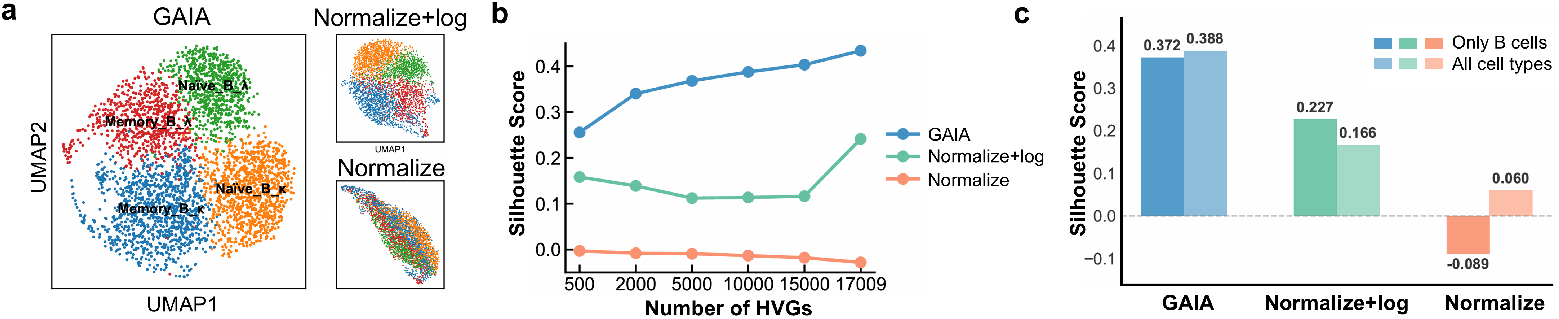
(a) GAIA identifies four B-cell subtypes. (b) SI scores of GAIA, Normalize, and Normalize+log when selecting different numbers of top HVGs. (c) Comparison of SI scores for the four B-cell subtypes under GAIA, Normalize, and Normalize+log using the full dataset and the B-cell–only subset.

To quantitatively assess the robustness and feature-independence of the GAIA classification, we computed the Silhouette score (SI) on the PCA embeddings obtained under each of the three frameworks, across multiple sets of highly variable genes (HVGs). Across all tested HVG configurations, the four GAIA-derived subtypes remained consistently well-separated, showing higher SI scores compared to normalization or normalization with log-transformation, as Fig 3b shows. This consistency indicates that GAIA captures intrinsic transcriptional geometry within the B cell population rather than being sensitive to arbitrary choices of input features. In other words, GAIA effectively extracts stable, geometry-preserving manifolds that are resilient to feature selection variations. Furthermore, when embedding B cells together with other major immune cell types, the same four GAIA-identified subtypes remained distinctly resolved within the global transcriptional landscape (Appendix A). The preservation of high SI scores (Fig 3c) in this broader context further demonstrates that GAIA achieves feature-independent, biologically reproducible delineation of B cell subtypes that generalizes beyond isolated clustering analyses. Collectively, these results establish GAIA as a robust and biologically grounded framework for subtyping immune cells, capable of maintaining consistent delineation across varying feature spaces and heterogeneous cellular environments.

As an additional validation, we applied GAIA to a human liver single-cell RNA-seq dataset (Appendix E) [26]. Consistent with the B cell analysis presented in the main text, GAIA achieved higher silhouette scores than normalization and normalization+log across different highly variable gene (HVG) selections. This supplementary analysis further supports the robustness of GAIA across datasets and feature choices. We also evaluated the computational efficiency of GAIA compared with standard normalization and normalization+log pipelines (Appendix F). As expected, GAIA incurs additional computational cost due to square-root mapping and tangent-space projection. In practice, the overall runtime scales linearly with the number of cells and genes and remains comparable to standard PCA-based workflows. Empirical benchmarks on datasets of comparable size indicate that GAIA incurs a moderate increase in preprocessing time, while remaining feasible for typical single-cell datasets. Detailed runtime and memory usage statistics are provided in Appendix F.

### 3.2 GAIA Enhances Spatial Domain Identification in Human DLPFC

Organic integration of qualitative and quantitative information is particularly important in spatial transcriptomics, where the diffusion of transcripts and the fusion of neighboring cells into spatial spots attenuate qualitative differences. We demonstrate the advantages of GAIA by benchmarking the domain segmentation performance of *scNiche* [27] on a human dorsolateral prefrontal cortex (DLPFC) dataset [28]. We compare annotations derived from GAIA-PCA, Euclidean PCA on normalized expression, and Euclidean PCA with normalization followed by log transformation. Fig 4 shows the domain segmentation results relative to manual annotation. Consistent with expectations, GAIA embeddings yield the most anatomically accurate domain delineations, which is reflected in a strong alignment with manually curated cortical layers and a high adjusted Rand index (ARI).

**Fig 4:**
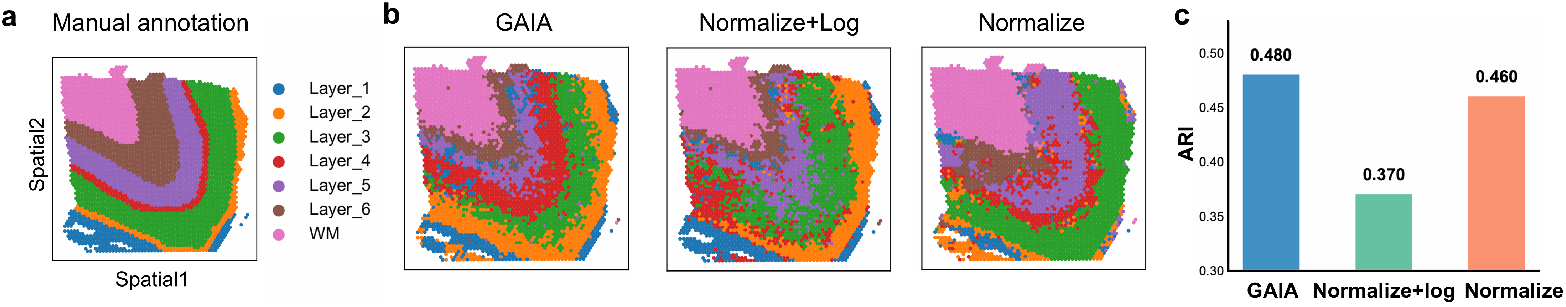
Domain detection results of the mouse dorsolateral prefrontal cortex layers with manual annotations using GAIA, normalized, and normalized+log embeddings via scNiche, along with ARI scores.

Notably, normalization followed by log transformation performs worst in domain segmentation. A primary contributor to its suboptimal performance is the attenuation of qualitative differences between cortical layers. Due to the limited spatial resolution of 10x Visium, each measured spot captures transcripts from multiple cells in a close vicinity. Transcriptomic differences between cortex layers manifest mostly as subtle quantitative shifts in average gene expression rather than clear presence/absence signals. Appendix B provides a detailed account of these effects. Together, these findings illustrate the benefit of harmonizing qualitative and quantitative variation with GAIA for spatial transcriptomics analyses.

### 3.3 GAIA Preserves Biological Structure Despite Sequencing Depth Variations

Sequencing-depth variation is a major source of batch effects in data integration. As sequencing depth decreases, moderately expressed genes progressively shift toward becoming high-dropout genes. When integrating datasets with drastically different depths, log-transformation amplifies inter-batch differences between low-dropout and high-dropout genes, creating spurious biological distinctions between batches.

To systematically evaluate the robustness of distance metrics against sequencing-depth variations, we benchmarked alignment of a public BMMC CITE-seq dataset against itself following random mRNA down-sampling [29]. Cell types were manually annotated based on surface marker expression [30]. We selected one BMMC dataset and randomly down-sampled mRNA molecules to 75%, 50%, and 25% of the original depth. Fig 5a displays the distribution of non-zero expression percentages across genes following down-sampling. As sequencing depth decreases, a substantial fraction of genes transition into high-dropout genes, accompanied by a corresponding decrease in non-zero expression cell percentages.

**Fig 5:**
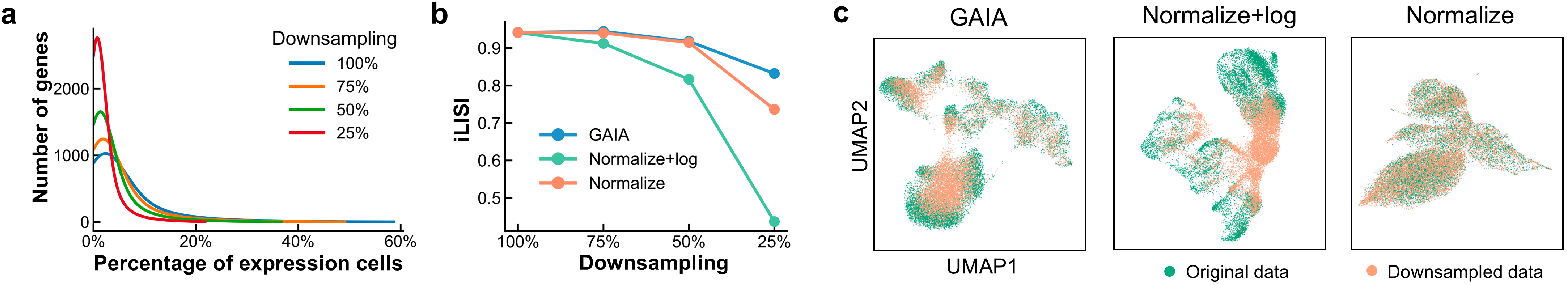
(a) Density plot of gene-expression percentages across downsampling rates. (b) iLISI scores for GAIA, normalize + log, and normalize at varying downsampling levels. (c) UMAP of GAIA, normalize + log, and normalize across downsampling rates.

When a functionally important gene transitions from a non-dropout to a high-dropout gene across datasets, log-transformation dramatically amplifies inter-batch differences. iLISI[31] scores, which quantify cross-batch cell intermixing, reveal that alignment degrades more rapidly under log-transformation than under GAIA or normalization-only approaches (Fig 5b). This degradation is visually apparent in UMAP plots(Fig 5c), where batch effects are most pronounced in the log-transformation framework.

GAIA uniquely maintains biological distinctions between cell types while minimizing batch mis-alignment. Compared to normalization-only approaches, GAIA achieves superior cell-type separation, reflected in higher cell-type silhouette indices (Appendix D). This advantage is evident in the four-batch integration results, where four BMMC samples are integrated into a common transcriptomic space without explicit batch correction. As Appendix D demonstrates, GAIA outperforms alternative methods by simultaneously delineating cell types and reducing batch effects. Collectively, these results demonstrate the superior performance of the GAIA dimension reduction and distance metric in multi-batch analysis.

## 4 Conclusion

In this study, we present GAIA, a framework that overcomes the limitations of conventional Euclidean or log-transformed embeddings by modeling cells as multinomial distributions and computing distances within an informational-geometric framework. This approach effectively balances quantitative and qualitative variations in gene expression, preserves robust cell-to-cell relationships, mitigates the batch effects from sequencing depth variations, and reduces reliance on feature selection. Notably, GAIA is applicable to both scRNA-seq and spatial transcriptomics, enhancing subtle transcriptomic differences, improving domain-level annotations, and providing a principled, interpretable, and depth-robust framework for single-cell and spatial analyses. **GAIA** is available at https://github.com/Carroll105/GAIA.

## 5 Acknowledgements

This work is supported by STI2030-Major Projects 2022ZD0212400, Lingang Laboratory grant LGL-8888, STCSM grant 24510714300 and 20DZ2254400, GuangDong Basic and Applied Basic Research Foundation grant 2023B1515120006 and SJTU Science-Medicine interdisciplinary grant YG2026ZD09 and 24X010301456. We also thank Prof. Hongyu Zhao for helpful discussions and insightful suggestions.

## References

1. Alev Baysoy, Zhiliang Bai, Rahul Satija, and Rong Fan. The technological landscape and applications of single-cell multi-omics. Nature Reviews Molecular Cell Biology, 24(10):695–713, 2023.

2. Yuge Ji, Tessa D Green, Stefan Peidli, Mojtaba Bahrami, Meiqi Liu, Luke Zappia, Karin Hrovatin, Chris Sander, and Fabian J Theis. Optimal distance metrics for single-cell rna-seq populations. BioRxiv, pages 2023–12, 2023.

3. Ebony Rose Watson, Ariane Mora, Atefeh Taherian Fard, and Jessica Cara Mar. How does the structure of data impact cell–cell similarity? evaluating how structural properties influence the performance of proximity metrics in single cell rna-seq data. Briefings in bioinformatics, 23(6):bbac387, 2022.

4. Taiyun Kim, Irene Rui Chen, Yingxin Lin, Andy Yi-Yang Wang, Jean Yee Hwa Yang, and Pengyi Yang. Impact of similarity metrics on single-cell rna-seq data clustering. Briefings in bioinformatics, 20(6):2316–2326, 2019.

5. F William Townes, Stephanie C Hicks, Martin J Aryee, and Rafael A Irizarry. Feature selection and dimension reduction for single-cell rna-seq based on a multinomial model. Genome biology, 20(1):295, 2019.

6. Pengyi Yang, Hao Huang, and Chunlei Liu. Feature selection revisited in the single-cell era. Genome Biology, 22(1):321, 2021.

7. Shaoheng Liang, Vakul Mohanty, Jinzhuang Dou, Qi Miao, Yuefan Huang, Muharrem Müftüoğlu, Li Ding, Weiyi Peng, and Ken Chen. Single-cell manifold-preserving feature selection for detecting rare cell populations. Nature computational science, 1(5):374–384, 2021.

8. Lucy Xia, Christy Lee, and Jingyi Jessica Li. Statistical method scdeed for detecting dubious 2d single-cell embeddings and optimizing t-sne and umap hyperparameters. Nature Communications, 15(1):1753, 2024.

9. Ian Covert, Rohan Gala, Tim Wang, Karel Svoboda, Uygar Sümbül, and Su-In Lee. Predictive and robust gene selection for spatial transcriptomics. Nature Communications, 14(1):2091, 2023.

10. Ziqi Rong, Jinpu Cai, Jiahao Qiu, Pengcheng Xu, Lana X Garmire, Qiuyu Lian, and Hongyi Xin. L2 normalization and geodesic distance for enhanced information preservation in visualizing high-dimensional single-cell sequencing data. In Proceedings of the 15th ACM International Conference on Bioinformatics, Computational Biology and Health Informatics, pages 1–11, 2024.

11. Constantin Ahlmann-Eltze and Wolfgang Huber. Comparison of transformations for single-cell rna-seq data. Nature Methods, 20(5):665–672, 2023.

12. Shun-ichi Amari. Information geometry and its applications, volume 194. Springer, 2016.

13. Henrique K Miyamoto, Fabio CC Meneghetti, Julianna Pinele, and Sueli IR Costa. On closed-form expressions for the fisher–rao distance. Information Geometry, 7(2):311–354, 2024.

14. Frank Nielsen. Fisher-rao and pullback hilbert cone distances on the multivariate gaussian manifold with applications to simplification and quantization of mixtures. In Topological, Algebraic and Geometric Learning Workshops 2023, pages 488–504. PMLR, 2023.

15. Min Cheol Kim, Rachel Gate, David S Lee, Andrew Tolopko, Andrew Lu, Erin Gordon, Eric Shifrut, Pablo E Garcia-Nieto, Alexander Marson, Vasilis Ntranos, et al. Method of moments framework for differential expression analysis of single-cell rna sequencing data. Cell, 187(22):6393–6410, 2024.

16. Yufeng Zhang, Jialu Pan, Li Ken Li, Wanwei Liu, Zhenbang Chen, Xinwang Liu, and Ji Wang. On the properties of kullback-leibler divergence between multivariate gaussian distributions. Advances in neural information processing systems, 36:58152–58165, 2023.

17. Sreangsu Acharyya, Arindam Banerjee, and Daniel Boley. Bregman divergences and triangle inequality. In Proceedings of the 2013 SIAM International Conference on Data Mining, pages 476–484. SIAM, 2013.

18. Victor M Panaretos and Yoav Zemel. Statistical aspects of wasserstein distances. Annual review of statistics and its application, 6(1):405–431, 2019.

19. Nihat Ay, Jürgen Jost, Hong Van Lê, and Lorenz Schwachhöfer. Information geometry, volume 64. Springer, 2017.

20. James G Dowty. Chentsov’s theorem for exponential families. Information geometry, 1(1):117–135, 2018.

21. Jason Osborne. Improving your data transformations: Applying the box-cox transformation. Practical Assessment, Research, and Evaluation, 15(1), 2010.

22. P Thomas Fletcher, Conglin Lu, Stephen M Pizer, and Sarang Joshi. Principal geodesic analysis for the study of nonlinear statistics of shape. IEEE transactions on medical imaging, 23(8):995–1005, 2004.

23. Tim Stuart, Andrew Butler, Paul Hoffman, Christoph Hafemeister, Efthymia Papalexi, William M Mauck III, Yuhan Hao, Marlon Stoeckius, Peter Smibert, and Rahul Satija. Comprehensive integration of single-cell data. Cell, 177(7):1888–1902, 2019.

24. Lisa M Rimsza, William A Day, Sarah McGinn, Anne Pedata, Yasodha Natkunam, Roger Warnke, James R Cook, Teresa Marafioti, and Thomas M Grogan. Kappa and lambda light chain mrna in situ hybridization compared to flow cytometry and immunohistochemistry in b cell lymphomas. Diagnostic pathology, 9(1):144, 2014.

25. Congxue Hu, Tengyue Li, Yingqi Xu, Xinxin Zhang, Feng Li, Jing Bai, Jing Chen, Wenqi Jiang, Kaiyue Yang, Qi Ou, et al. Cellmarker 2.0: an updated database of manually curated cell markers in human/mouse and web tools based on scrna-seq data. Nucleic acids research, 51(D1):D870–D876, 2023.

26. Benjamin J Stewart, John R Ferdinand, Matthew D Young, Thomas J Mitchell, Kevin W Loudon, Alexandra M Riding, Nathan Richoz, Gordon L Frazer, Joy UL Staniforth, Felipe A Vieira Braga, et al. Spatiotemporal immune zonation of the human kidney. Science, 365(6460):1461–1466, 2019.

27. Jingyang Qian, Xin Shao, Hudong Bao, Yin Fang, Wenbo Guo, Chengyu Li, Anyao Li, Hua Hua, and Xiaohui Fan. Identification and characterization of cell niches in tissue from spatial omics data at single-cell resolution. Nature Communications, 16(1):1693, 2025.

28. Kristen R Maynard, Leonardo Collado-Torres, Lukas M Weber, Cedric Uytingco, Brianna K Barry, Stephen R Williams, Joseph L Catallini, Matthew N Tran, Zachary Besich, Madhavi Tippani, et al. Transcriptome-scale spatial gene expression in the human dorsolateral prefrontal cortex. Nature neuroscience, 24(3):425–436, 2021.

29. Christopher Lance, Malte D Luecken, Daniel B Burkhardt, Robrecht Cannoodt, Pia Rautenstrauch, Anna Laddach, Aidyn Ubingazhibov, Zhi-Jie Cao, Kaiwen Deng, Sumeer Khan, et al. Multimodal single cell data integration challenge: results and lessons learned. BioRxiv, pages 2022–04, 2022.

30. Marlon Stoeckius, Christoph Hafemeister, William Stephenson, Brian Houck-Loomis, Pratip K Chattopadhyay, Harold Swerdlow, Rahul Satija, and Peter Smibert. Simultaneous epitope and transcriptome measurement in single cells. Nature methods, 14(9):865–868, 2017.

31. Ilya Korsunsky, Nghia Millard, Jean Fan, Kamil Slowikowski, Fan Zhang, Kevin Wei, Yuriy Baglaenko, Michael Brenner, Po-ru Loh, and Soumya Raychaudhuri. Fast, sensitive and accurate integration of singlecell data with harmony. Nature methods, 16(12):1289–1296, 2019.

