## supplementary for "Information Geometry Reconciles Discrete and Continuous Variation in Single-Cell and Spatial Transcriptomic Analysis"

#### Appendix A:GAIA harmonizes qualitative and quantitative differences in gene expression

A central challenge in single-cell data normalization is maintaining a balance between qualitative (discrete, on/off marker expression) and quantitative (continuous, graded expression) variation. Standard transformations, such as the log or square-root functions, often bias one aspect over the other—amplifying low-expression noise while compressing high-expression dynamics. To systematically explore how transformation choices affect this qualitative–quantitative balance, we adopted the Box–Cox transformation as a unifying analytical framework. Its tunable parameter

$\lambda$ allows a continuous interpolation between linear and logarithmic scaling, providing a principled way to examine how different levels of nonlinearity influence biological signal preservation.

The Box–Cox transformation is a flexible family of power transformations commonly used to stabilize variance and normalize data distributions. It is defined as

It is defined as

$$y(\lambda) = \begin{cases} \frac{x^\lambda - 1}{\lambda}, & \lambda \neq 0, \\ \log(x), & \lambda = 0, \end{cases} \tag{1}$$

where $\lambda$ is a tunable parameter controlling the degree of transformation. By adjusting $\lambda$, the Box-Cox transformation provides a continuum from no transformation ($\lambda = 1$) to log transformation ($\lambda = 0$), allowing gene expression data to be rescaled in a manner that can either emphasize or compress high-expression genes relative to low-expression genes.

To evaluate how different transformations affect the preservation of both qualitative and quantitative variation, we simulated a dataset containing three cell types. Here, qualitative differences are measured by the distance between the class centers of cell types A and B, reflecting mutually exclusive or on/off gene expression programs, while quantitative differences are measured by the distance between the class centers of cell types A and C, representing graded changes in gene expression. Using class-center distances provides an interpretable metric of biologically meaningful variation.

We then defined a *harmonization score* to assess how well a transformation balances these two types of differences. Specifically, if $D_{\text{qual}}$ denotes the class-center distance for qualitative differences (A vs. B) and $D_{\text{quant}}$ denotes the class-center distance for quantitative differences (A vs. C), the harmonization score $H$ is computed as their harmonic mean:

$$H = \frac{2}{\frac{1}{D_{\text{qual}}} + \frac{1}{D_{\text{quant}}}}. \tag{2}$$

By taking the harmonic mean, $H$ penalizes transformations that preserve one type of difference at the expense of the other, ensuring that both qualitative and quantitative separations are maintained.

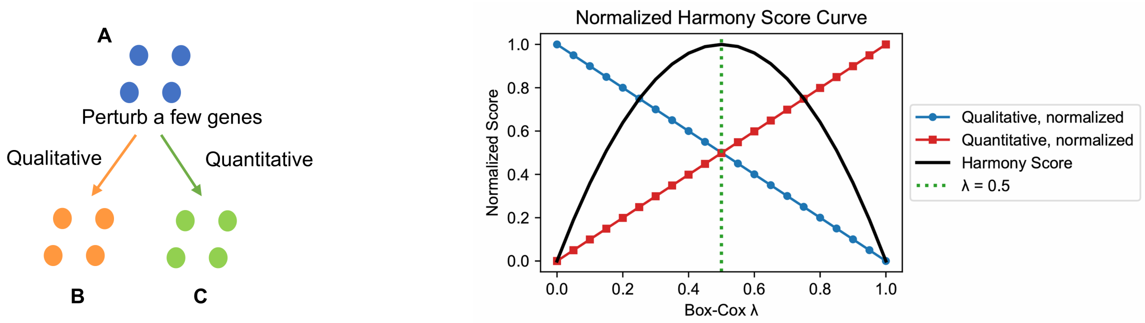


Fig. 1: Harmonization score of simulated data under Box-Cox transformation

Systematic evaluation across different Box-Cox transformations revealed that the harmonization score peaks at $\lambda = 0.5$ as Fig1 shows, corresponding to the square-root transformation. This indicates that the square-root transformation optimally balances the preservation of both mutually exclusive low-expression marker genes and graded high-expression signals. In contrast, no transformation ($\lambda = 1$) overemphasizes high-expression genes, while log transformation ($\lambda = 0$) exaggerates low-expression noise, compressing quantitative differences.

These results suggest that the square-root transformation is a principled choice for harmonizing both types of variation, enabling downstream analyses—including clustering, trajectory inference, and differential expression—to effectively capture both graded and binary gene expression patterns.

### Appendix B:GAIA Robustly Identifies B Cell Subtypes Across Feature Selection and Heterogeneous Contexts

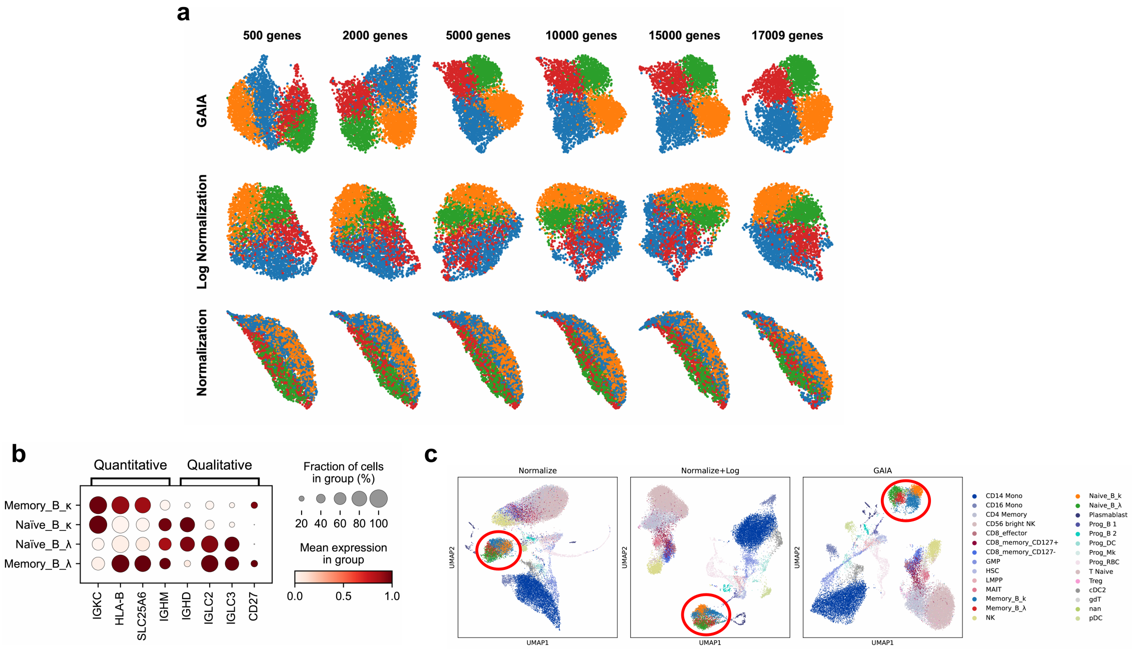


Fig. 2: GAIA identifies robust B cell subtypes under varying conditions. (a) UMAP visualization of B cells with different numbers of highly variable genes (HVGs) selected, showing that GAIA-derived embeddings consistently preserve the separation of the four B cell subtypes. (b) Dot plot of marker genes across the four B cell clusters. Both quantitative (gradual changes in expression levels) and qualitative (on/off presence) differences are captured, demonstrating the ability of GAIA to retain multiple aspects of transcriptional variation. (c) UMAP of GAIA embeddings for B cells integrated within all major BMMC cell types, showing that the four B cell subtypes remain distinctly separated in the manifold, even in the context of heterogeneous cell populations.

Using GAIA embeddings, B cells were robustly separated into four distinct subtypes across different feature selection settings. UMAP visualization (Fig2a) showed that the four clusters remained clearly separated regardless of the number of highly variable genes (HVGs) used, indicating that GAIA captures intrinsic transcriptional structures without heavy reliance on specific feature sets. Marker gene analysis revealed that these clusters were enriched for known B cell markers, with dot plots (Fig2b) highlighting both quantitative differences (gradual changes in expression levels) and qualitative differences (on/off expression), demonstrating that GAIA preserves multiple aspects of transcriptional variation. When embedding B cells within all major BMMC cell types, GAIA maintained clear separation of the four B cell subtypes on the manifold (Fig2c), indicating that the identified subtypes are robust even in the context of heterogeneous cell populations.

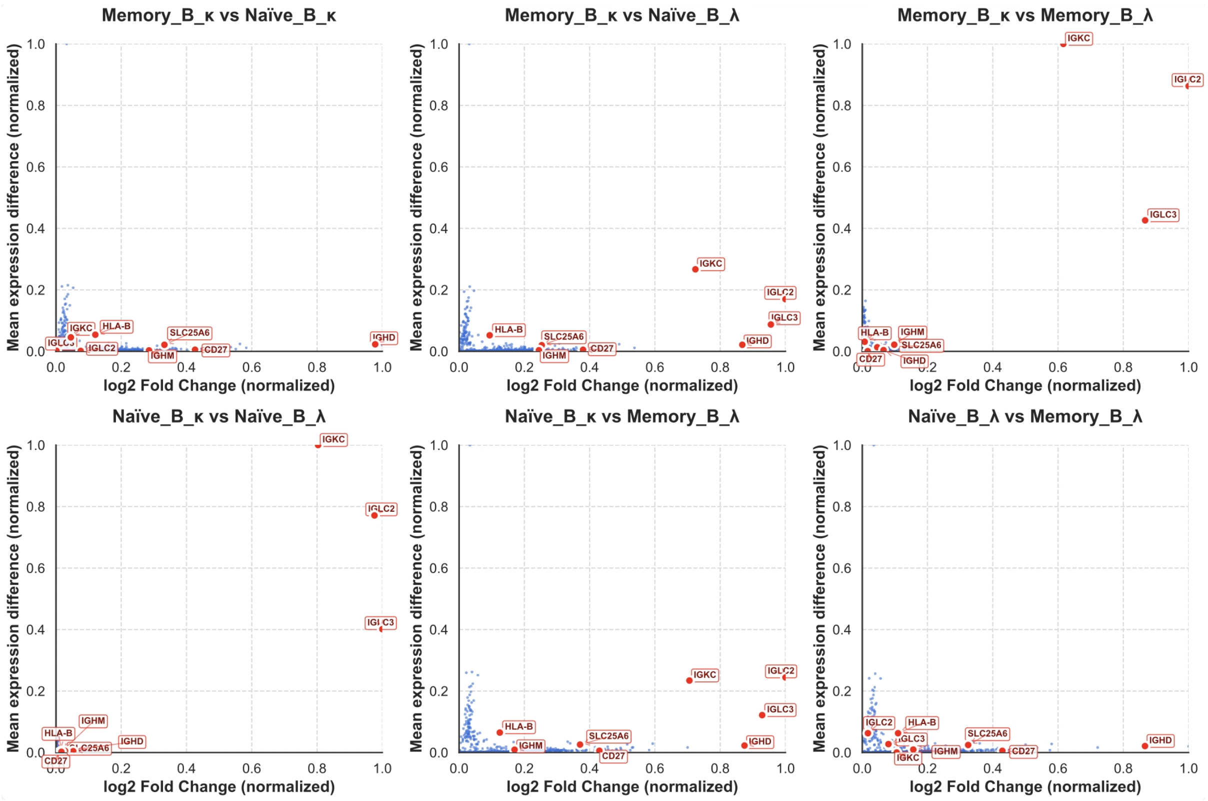


Fig. 3: The scatter plot showing the mean expression difference and log fold change (FC) between the four B cell subtypes. Both values were min–max normalized to the range of 0–1 for visualization purposes.

To further dissect these two aspects, we computed for each gene the difference in mean expression and the log2 fold change between every pair of B cell subtypes (Fig3). These two values respectively capture quantitative and qualitative expression changes, allowing us to classify genes according to whether their variation across subtypes is primarily quantitative or qualitative. Interestingly, many genes exhibited mixed behaviors—showing qualitative differences between some subtype pairs but quantitative shifts between others—suggesting that a balance between quantitative and qualitative transcriptional variation is crucial for defining B cell identities. Together, these results demonstrate that GAIA provides a stable and feature-independent framework for B cell subtype identification.

### Appendix C: GAIA Preserves Layer-Specific Differences and Robust Neighborhood Structures in Cortical Domains

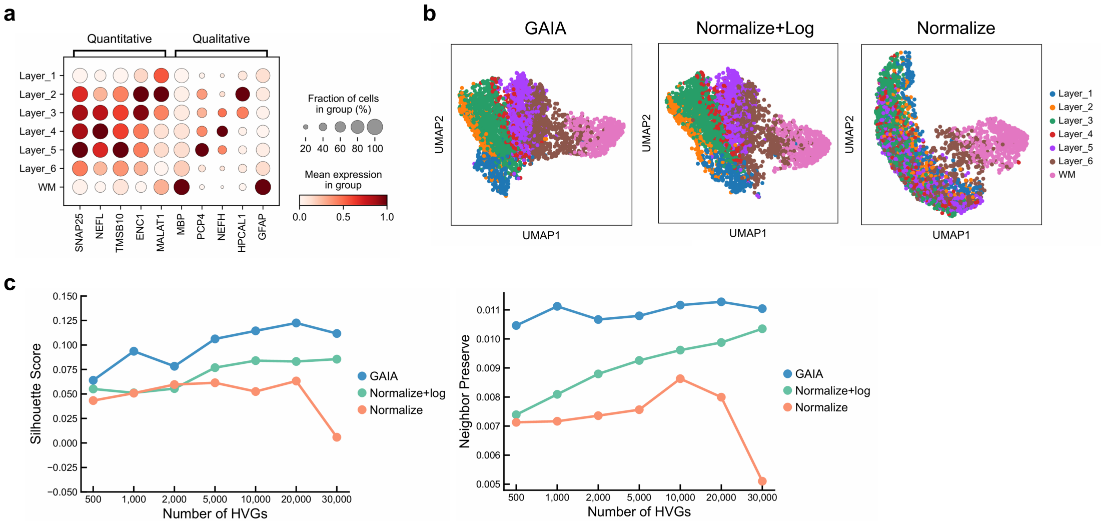


Fig. 4: GAIA preserves layer-specific transcriptional differences and robust neighborhood structures. (a) Expression of domain marker genes across cortical layers, highlighting that differentially expressed genes between domains exhibit both qualitative (on/off) and quantitative (gradual) differences. (b) UMAP embeddings of cortical layers under different transformations. GAIA-derived embeddings most effectively preserve the transcriptional differences between layers compared to standard normalization or log-transformed embeddings. (c) Comparison of Separation Index (SI) scores and Neighbor Preservation scores across different feature selection sizes, demonstrating the robustness of GAIA in maintaining layer separations and local neighborhood structures.

To evaluate the performance of GAIA in capturing layer-specific transcriptional variation, we examined domain marker gene expression, UMAP embeddings, and clustering metrics. Marker analysis (Fig5a) revealed that differentially expressed genes between domains contained both quantitative and qualitative differences, reflecting subtle and discrete transcriptional variations. UMAP visualization under different transformations (Fig5b) showed that GAIA embeddings most effectively preserved these layer-specific differences, maintaining clear separations between cortical layers. Quantitative assessment using SI scores and Neighbor Preservation scores (Fig5c) confirmed that GAIA consistently outperformed standard normalization and log-transformed embeddings across varying numbers of selected features, indicating that GAIA provides a robust, feature-independent framework for domain-level spatial mapping.

### Appendix D: GAIA effectively preserves biologically meaningful structure and remains insensitive to changes in sequencing depth

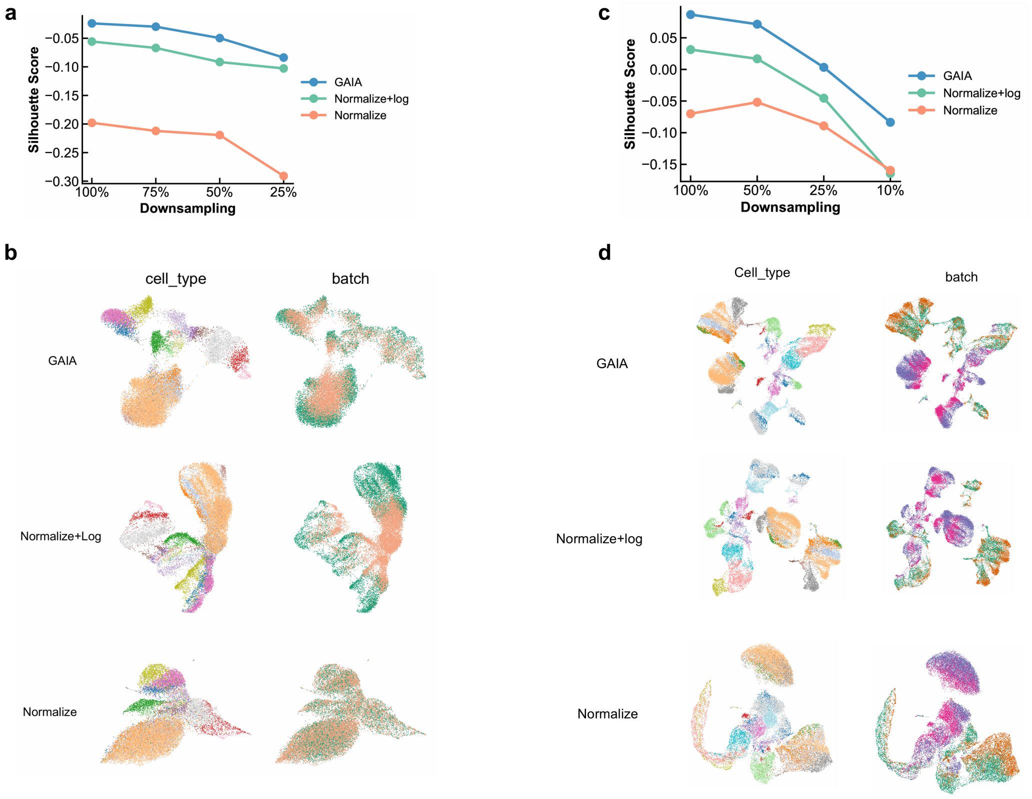


Fig. 5: (a) SI scores for Normalize, Normalize + log, and GAIA computed at different downsampling rates for a single donor. For each downsampling rate, SI scores were calculated using the original dataset and its downsampled counterpart. (b) SI scores for Normalize, Normalize + log, and GAIA across four donors at multiple downsampling rates. Each donor was independently downsampled at several rates, and SI scores were computed based on the resulting downsampled datasets. (c) Corresponding results for the single donor in (a): UMAP plots of Normalize, Normalize + log, and GAIA, colored by cell type and by batch. (d) Corresponding results for the four donors in (b): UMAP plots of Normalize, Normalize + log, and GAIA, colored by cell type and by batch.

### Appendix E: GAIA preserves cluster integrity in human liver data

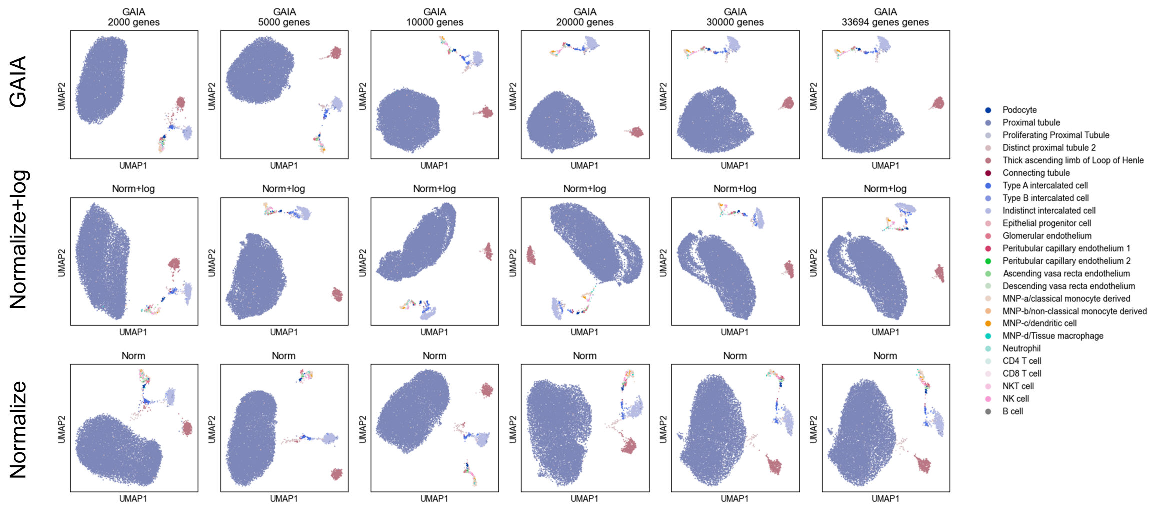


Fig. 6: UMAP of human liver cells comparing standard normalization, log-normalization, and GAIA.

To assess GAIA's performance on human liver single-cell RNA-seq data, we compared it with standard normalization and log-transformation under varying feature selection settings. We first evaluated cluster quality using silhouette (SI) scores across different feature selections. GAIA consistently achieved the highest SI values (Figure 7), indicating superior preservation of cluster integrity compared with the other methods. Visual inspection of proximal tubule cells further supported this observation (Figure6). While normalization + log appeared to separate these cells into two subclusters, both GAIA and standard normalization maintained the coherent structure of all clusters, regardless of the number of selected genes. This suggests that the apparent subcluster identified under normalization + log is likely an artifact rather than a biologically meaningful subtype.

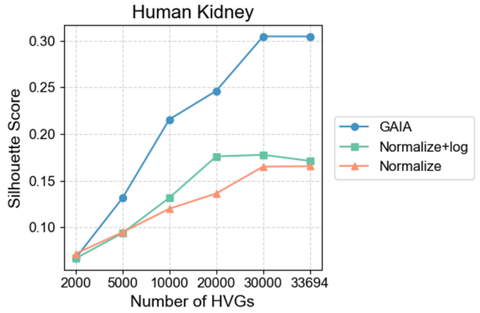


Fig. 7: SI score of human liver cells comparing standard normalization, log-normalization, and GAIA.

### Appendix F: Runtime analysis

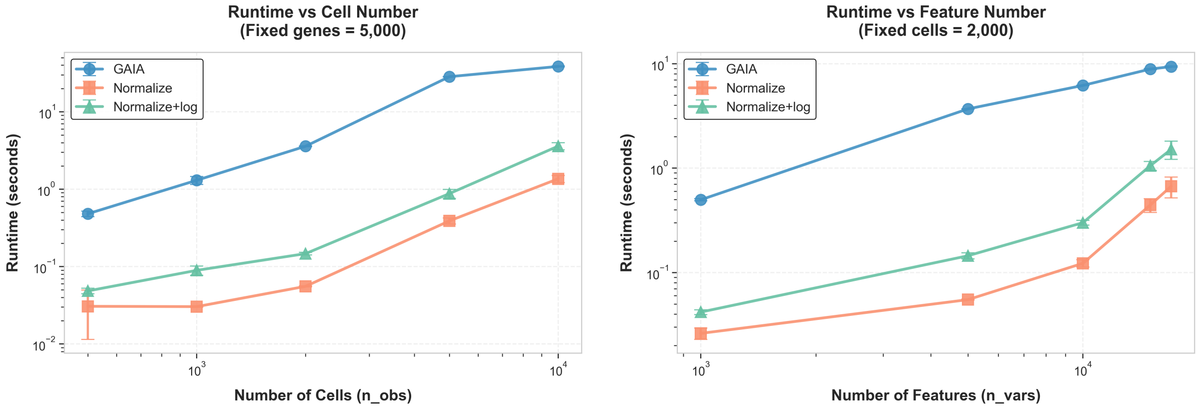


Fig. 8: Runtime comparison of three preprocessing methods. Left: Runtime versus cell number ($n_{\text{obs}}$; 5,000 genes fixed). Right: Runtime versus feature number ($n_{\text{vars}}$; 2,000 cells fixed). Methods include GAIA, Normalize, and Normalize+log. Error bars show standard deviation across three runs, and both axes are log-scaled. GAIA exhibits moderate but practical computational overhead compared with standard pipelines.

We compared the computational efficiency of three preprocessing pipelines: GAIA, Normalize (library size normalization + PCA), and Normalize+log (normalization + log-transformation + PCA). We evaluated scalability by varying (i) the number of cells (500–10,000; 5,000 genes fixed) and (ii) the number of features (1,000–17,000; 2,000 cells fixed). Each configuration was repeated three times, extracting 40 principal components.

Across all settings, the runtime ranking was consistent: Normalize was fastest, Normalize+log showed moderate overhead due to log-transformation, and GAIA was the slowest, because of square-root mapping and tangent space projection. Despite this overhead, GAIA remains practical for standard single-cell datasets; even for 10,000 cells and 5,000 genes, runtime is approximately 40 seconds. Given that preprocessing is performed once per dataset, the moderate computational cost is justified by improved biological signal preservation.
